# Robust Genome-Wide Ancestry Inference for Heterogeneous Datasets and Ancestry Facial Imaging based on the 1000 Genomes Project

**DOI:** 10.1101/549881

**Authors:** Jairui Li, Tomas Gonzalez, Julie D. White, Karlijne Indencleef, Hanne Hoskens, Alejandra Ortega Castrillon, Nele Nauwelaers, Arslan Zaidi, Ryan J. Eller, Torsten Günther, Emma M. Svensson, Mattias Jakobsson, Susan Walsh, Kristel Van Steen, Mark D. Shriver, Peter Claes

## Abstract

Accurate inference of genomic ancestry is critically important in human genetics, epidemiology, and related fields. Geneticists today have access to multiple heterogeneous population-based datasets from studies collected under different protocols. Therefore, joint analyses of these datasets require robust and consistent inference of ancestry, where a common strategy is to yield an ancestry space generated by a reference dataset. However, such a strategy is sensitive to batch artefacts introduced by different protocols. In this work, we propose a novel robust genome-wide ancestry inference method; referred to as SUGIBS, based on an unnormalized genomic (UG) relationship matrix whose spectral (S) decomposition is generalized by an Identity-by-State (IBS) similarity degree matrix. SUGIBS robustly constructs an ancestry space from a single reference dataset, and provides a robust projection of new samples, from different studies. In experiments and simulations, we show that, SUGIBS is robust against individual outliers and batch artifacts introduced by different genotyping protocols. The performance of SUGIBS is equivalent to the widely used principal component analysis (PCA) on normalized genotype data in revealing the underlying structure of an admixed population and in adjusting for false positive findings in a case-control admixed GWAS. We applied SUGIBS on the 1000 Genome project, as a reference, in combination with a large heterogeneous dataset containing auxiliary 3D facial images, to predict population stratified average or ancestry faces. In addition, we projected eight ancient DNA profiles into the 1000 Genome ancestry space and reconstructed their ancestry face. Based on the visually strong and recognizable human facial phenotype, comprehensive facial illustrations of the populations embedded in the 1000 Genome project are provided. Furthermore, ancestry facial imaging has important applications in personalized and precision medicine along with forensic and archeological DNA phenotyping.

**Author Summary:** Estimates of individual-level genomic ancestry are routinely used in human genetics, epidemiology, and related fields. The analysis of population structure and genomic ancestry can yield significant insights in terms of modern and ancient population dynamics, allowing us to address questions regarding the timing of the admixture events, and the numbers and identities of the parental source populations. Unrecognized or cryptic population structure is also an important confounder to correct for in genome-wide association studies (GWAS). However, to date, it remains challenging to work with heterogeneous datasets from multiple studies collected by different laboratories with diverse genotyping and imputation protocols. This work presents a new approach and an accompanying open-source software toolbox that facilitates a robust integrative analysis for population structure and genomic ancestry estimates for heterogeneous datasets. Given that visually evident and easily recognizable patterns of human facial characteristics covary with genomic ancestry, we can generate predicted ancestry faces on both the population and individual levels as we illustrate for the 26 1000 Genome populations and for eight eminent ancient-DNA profiles, respectively.

## Introduction

Scientists today have access to large heterogeneous datasets from many studies collected by different laboratories with diverse genotyping and imputation protocols. The joint analysis of these datasets requires a robust and consistent inference of ancestry across all datasets involved, where one common strategy is to yield an ancestry space generated by a reference set of individuals (1). Based on open-research initiatives such as the 1000 Genome project (1KGP) (2), HapMap project (3), Human Genome Diversity project (HGDP) (4), and the POPRES dataset (5), the potential exists to create reference ancestry latent-spaces at different levels of interest, from worldwide inter-continental to fine-scale intra-continental ancestry. A reference ancestry space allows the researcher to collate multiple datasets facilitating analyses that are more advanced. For example, reference ancestry spaces can be used to infer the population structure of samples with family structure or cryptic relatedness (1) and to investigate the genetic similarity between ancient DNA and modern human genomes (6). They also have the potential to correct for population structure in a genome-wide association study (GWAS) on heterogeneous and admixed samples. Of final interest is the association of auxiliary data (e.g. specific phenotypes, such as 3D facial shape used in this work) present in internally collected datasets with ancestral variations captured by a reference space. This requires the projection of the collected datasets into a reference space, followed by an association of the projection scores with the auxiliary data presented.

Methodologically, the idea is to construct an ancestry latent-space from a reference dataset and to enable the projection of new cases from other datasets that follow the mainstream of the reference dataset. Starting from genome-wide single nucleotide polymorphisms (SNPs), PCA and analogous dimension reduction techniques on normalized genotype data are popular strategies used in this context (7,8). However, in construction of an ancestry space, these approaches are known to be sensitive to outliers (7,9). In addition and more importantly, in projecting new cases onto an ancestry space, PCA produces patterns of misalignment (for example, “shrinkage” patterns where projected cases tend to falsely gravitate towards the center of the ancestry space) due to missing data, missing heterozygotes, and genotyping along with imputation errors, which is misleading without careful interpretation (1). Therefore, stringent quality control and data filters are typically in place to remove individual outliers and SNP data with high missing rates or not in Hardy-Weinberg equilibrium (HWE). However, in heterogeneous datasets, in contrast to homogeneous datasets, such data filters are harder to define, and potentially remove SNP data related to population structure. Furthermore, genotyping and imputation batch artefacts, not detected by quality control and different from one protocol to another, typically remain and still affect an integrative analysis of ancestry.

In this work, we propose a novel robust genome-wide ancestry inference (referred to as SUGIBS) based on the spectral (S) decomposition of an unnormalized genomic (UG) relationship matrix generalized by an Identity-by-State (IBS) similarity degree of individuals’ matrix. Robustness against outliers, during ancestry space construction, is obtained by absence of specific sample statistics (e.g. allele frequencies). Furthermore, SUGIBS provides a robust projection of new samples, from different studies, onto a reference SUGIBS space. During projection, the IBS similarity degree of individuals to project to individuals in the reference dataset acts as a correcting term for missing genotypes and errors, and most interestingly this correction is on an individual-by-individual basis. We test the robustness of SUGIBS and compare its performance to PCA and Multi-Dimensional Scaling (MDS) in revealing the underlying structure of an admixed population and adjusting for false positive findings in a simulated case-control admixed GWAS. Using the 1KGP as reference dataset, and an additional heterogeneous dataset containing 3D facial images, we apply SUGIBS to construct ancestry faces that illustrate the ancestral variation captured in the 1KGP. Additionally, we reconstruct the ancestry faces for eight high-coverage ancient DNA genomes further illustrating the potential of the work. Based on the results, our method facilitates a robust integrative analysis for ancestry estimation in heterogeneous datasets.

## Results

In the first experiment, we investigated the robustness of SUGIBS in comparison to traditional approaches, in particular PCA using normalized or unnormalized genotype data and MDS using IBS distances as they are implemented in PLINK 1.9 (10), against individual outliers in a reference dataset. For this purpose, we first selected all unrelated individuals from the CEU and TSI populations in the HapMap 3 project (Belmont et al., 2003) and used SUGIBS, PCA, unnormalized PCA (UPCA) and MDS to illustrate the first and second latent dimensions as ancestry components (Figure 1, top row). In contrast to the traditionally used normalized genotypes in PCA, UPCA used unnormalized genotypes that were not centralized around the mean and were not standardized to a variance equal to one. As expected, PCA, MDS and SUGIBS are able to differentiate between both populations along the first ancestry component. The first component of UPCA seems to aggregate the average pattern of SNPs instead of the differentiation between two groups. Surprisingly, with PCA a single outlier (NA11917) that was not expected during the selection of both populations already affected the second ancestry component. Subsequently, we randomly selected one individual from four different and additional populations (CHB, GIH, MEX and YRI) as “outliers” in the dataset. Figure 1, bottom row, illustrates the first two ancestry components of the four methods constructed on the dataset with outliers, where all four approaches clearly separate the outliers. Using PCA, in contrast to MDS, UPCA and SUGIBS the clear distinction between CEU and TSI is lost within the first two ancestry components, as they mainly capture variations due to the outliers. The main reason for robustness in UPCA, MDS and SUGIBS is that these three methods use unnormalized genotype data and therefore do not rely on specific sample statistics (e.g. allele frequencies), that otherwise increase the influence of outlier variation.

**Figure 1:**
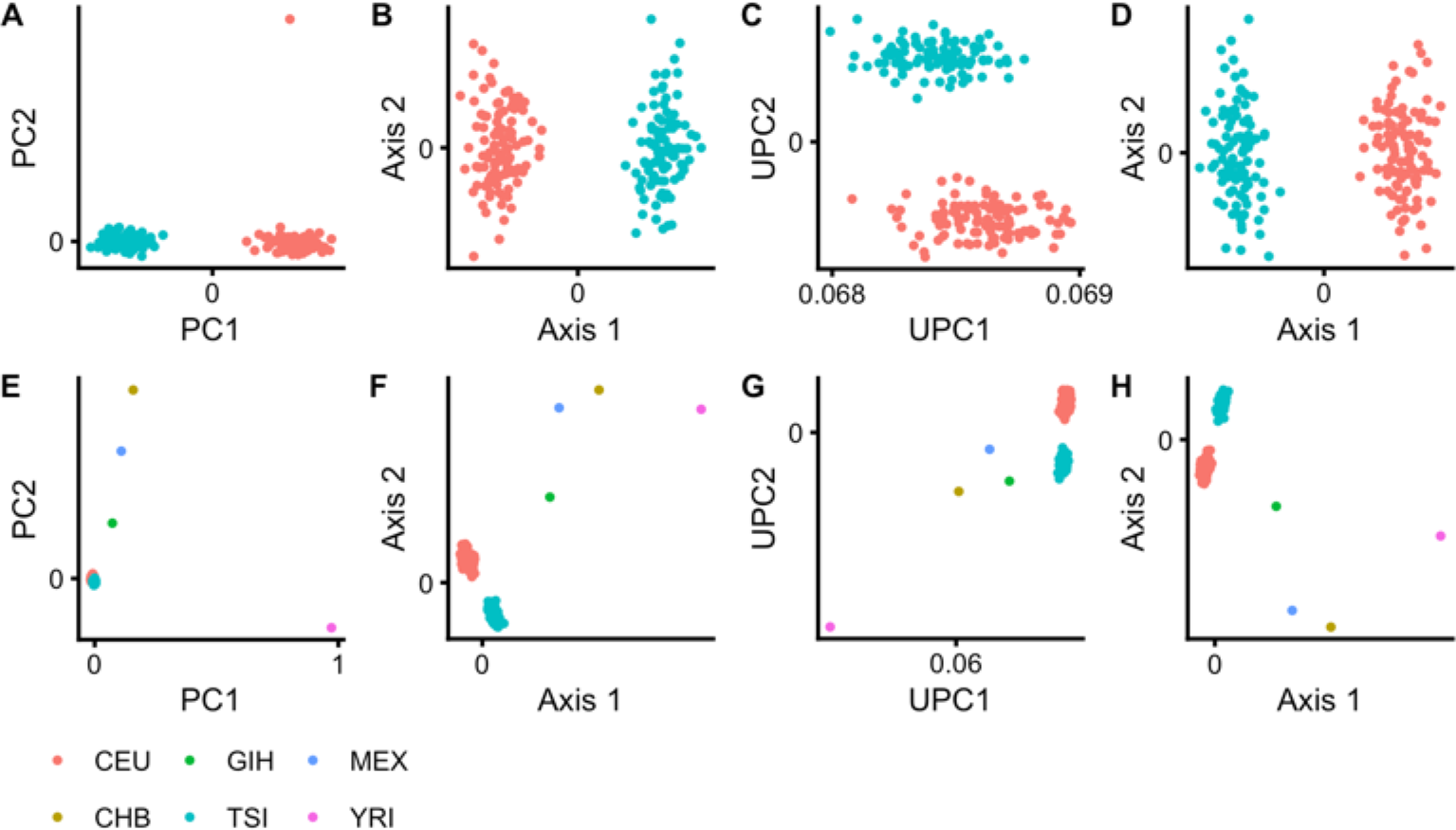
Robustness against individual outliers during the construction of an ancestry space. Top row, the first two ancestry components for A) PCA, B) MDS, C) UPCA and D) SUGIBS using the CEU and TSI populations from the HapMap 3 project. Bottom row, the first two ancestry components for E) PCA, F) MDS, G) UPCA and H) SUGIBS using the CEU and TSI populations from the HapMap 3 project, but with randomly selected single individuals from four different and additional populations (CHB, GIH, MEX and YRI) as “outliers”.

In a second experiment, we projected (Methods, equation 4) new samples on an ancestry space, based on the 1KGP as reference dataset, to investigate the robustness of SUGIBS in comparison to PCA and UPCA against typical artifacts of different laboratory protocols. Note that, since the first component of UPCA just aggregated the average pattern as seen in experiment 1, we started UPCA from the second component onwards. Also note that, MDS does not allow for a straightforward projection of new samples on a reference space and was therefore excluded. As samples to project, we randomly assigned all 1,043 individuals of 51 populations from the HGDP dataset (4) into two equally-sized samples, one unchanged and one modified, respectively. To investigate the influence of different rates of missing data, we randomly masked 5% of the SNP genotypes as missing in the modified population (See Methods). For the influence of different rates of errors, we partially changed SNP genotypes with minor allele frequency (MAF) less than 5% in the modified population (See Methods). Note that this was done knowing that more imputation errors are observed in SNPs with a MAF of 5% and less (11). We projected both HGDP populations onto the PCA, UPCA and SUGIBS reference spaces as defined by the 1KGP. In PCA, the simulated artefacts generated “shrinkage” and “shifting” patterns of misalignment in the first two projected ancestry components (Figure 2, top row), for missing and erroneous genotypes, respectively. UPCA was only influenced by missing genotypes (Figure 2, middle row). In contrast, SUGIBS was not influenced by missing or erroneous genotypes (Figure 2, bottom row). Figure 3 summarizes the normalized root-mean-square deviations (NRMSD) of the first eight axes of SUGIBS, UPCA and PCA of the modified HGDP population over 100 simulations. SUGIBS is significantly more robust than PCA in the presence of missing and genotyping/imputation errors in new data for which ancestry needs to be inferred, by projecting it into a reference space.

**Figure 2:**
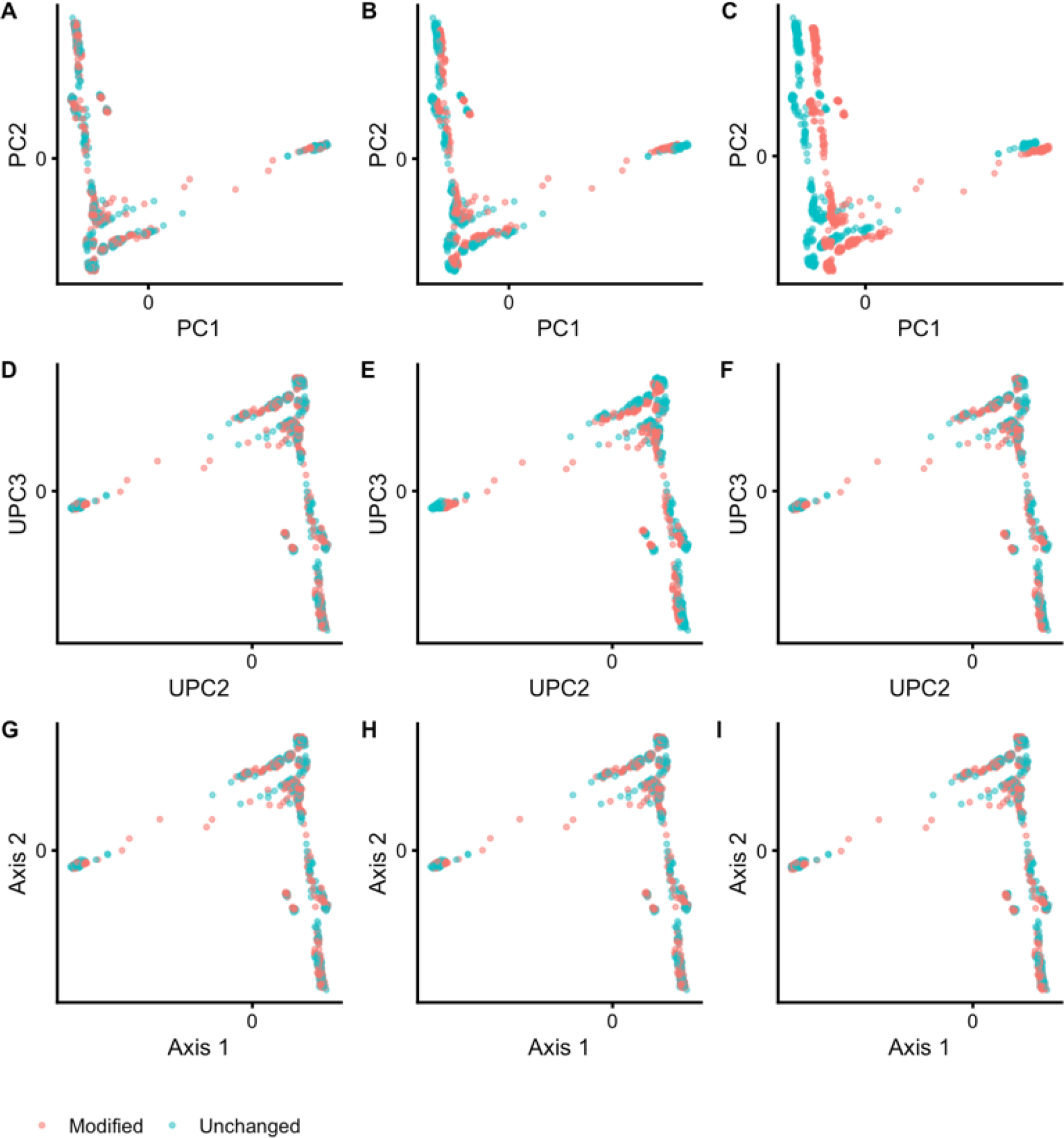
Robustness against batch artefacts during the projection of samples onto an ancestry space. Top row, the first two ancestry components of PCA using the original genotypes A), missing genotypes B) and modified genotypes C). Middle row, the second and third ancestry components of UPCA using the original genotypes D), missing genotypes E) and modified genotypes F). Bottom row, the first two ancestry components of SUGIBS using the original genotypes G), missing genotypes H) and modified genotypes I).

**Figure 3:**
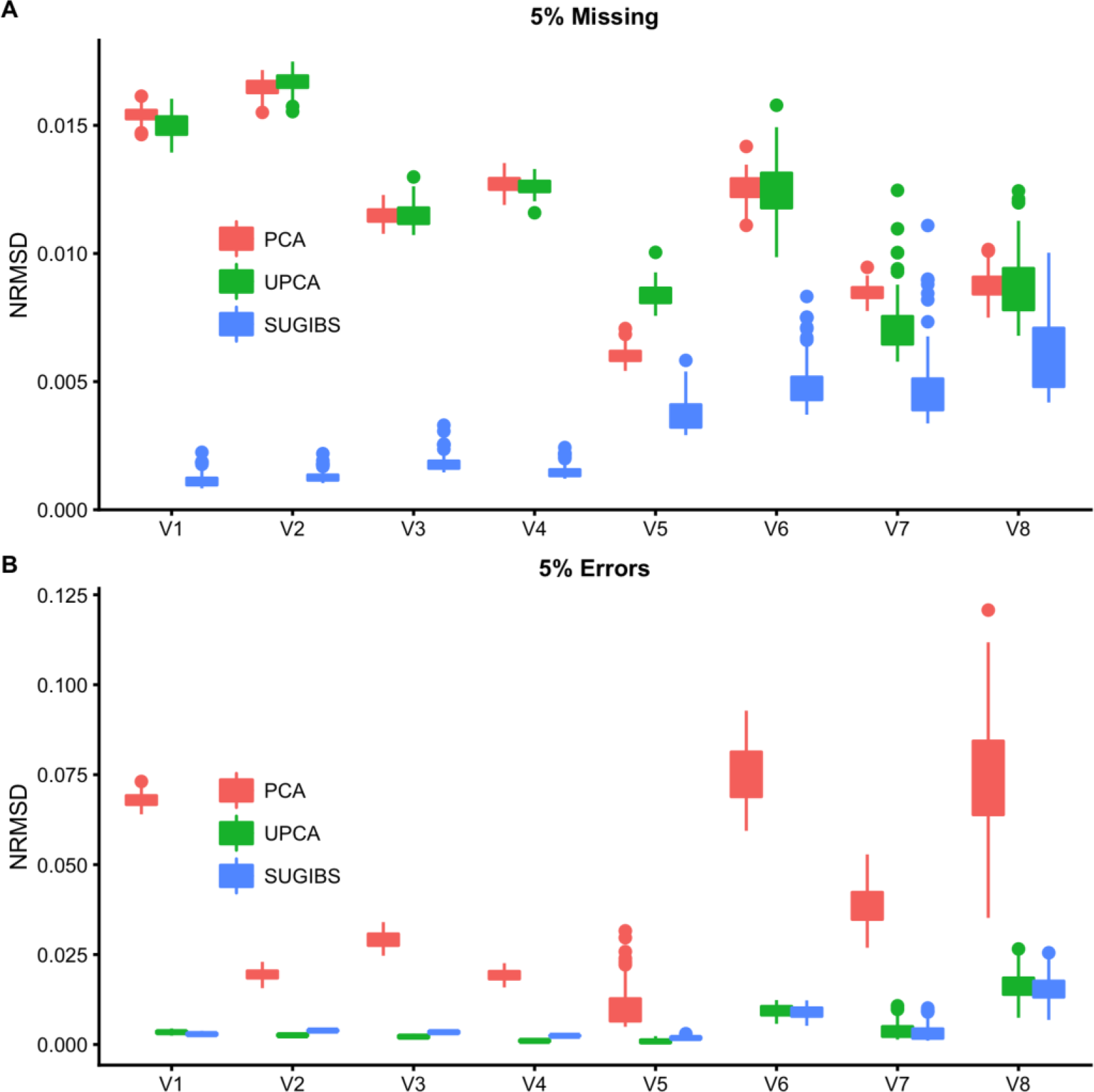
Normalized root-mean-square deviation (NRMSD) of the top eight axes of PCA, UPCA and SUGIBS. NRMSD measures the root-mean-square differences (RMSD), for the modified HGDP population only between the scores on ancestry axes generated using the original genotypes (error free) and the modified genotypes (with simulated errors, A) missing genotypes and B) erroneous genotypes). The RMSD values were normalized by the range of the ancestry axes generated using the original genotypes, so that NRMSD of the three methods (PCA, UPCA and SUGIBS) are comparable.

In a third experiment, following the work of Galinsky et al. (12), we investigated the ability of SUGIBS compared to PCA and MDS in representing admixture. We simulated data at 10,000 random independent SNPs for 1,000 individuals from a recent admixture of two populations, 50% from each population on average with divergences *F*_*st*_ = {0.001,0.005,0.01,0.05,0.1}, from an intra-European difference to an intercontinental difference (13). Because the admixture contains only one dimension of population structure, only the first component of variation is of interest. Figure 4 presents the absolute correlations between the first component of PCA, MDS and SUGIBS and the simulated ancestry proportions over 100 runs. When the *F*_*st*_ divergence between two populations is lower than 0.05, the correlation between the SUGIBS component and the ancestry proportion is similar to that of MDS, but a little lower than PCA. We noticed that when *F*_*st*_ ≤ 0.01, all three methods have a reduced performance to reveal the underlying admixture and when *F*_*st*_ > 0.01, all three methods perform perfectly.

**Figure 4:**
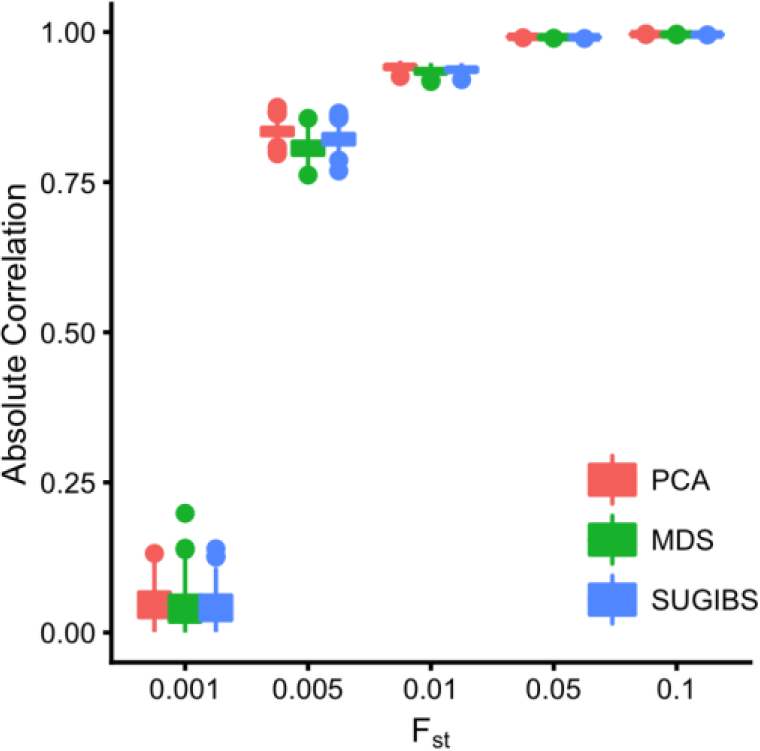
Capturing simulated admixture in function of F_st_. X-axis represents the different levels of Fst investigated. The Y-axis represents the absolute correlation of the first component in PCA, MDS and Spectral-IBS with the simulated ancestry proportion. The higher the correlation the better a method is able to capture the underlying admixture.

Following the work of Price et al. (14), we also simulated a case-control GWAS to investigate if the population structure inferred by SUGIBS can be used for correcting population stratification as a confounder. Only low divergences between the two populations *F*_*st*_ = {0.001,0.005,0.01}, were tested, because for larger divergences all three methods would perform the same as deducted from the previous experiment. Tests were conducted with a logistic regression under four different correction scenarios: 1) no population for stratification correction (Naïve), 2) PCA, 3) MDS and 4) SUGIBS, using a likelihood ratio test for the significance of each genetic marker. The experiment was conducted 100 times, with average proportions of SNPs detected as significant shown in Table 1. These results indicate that in a single dimensional population structure, correcting using MDS, SUGIBS and PCA perform similarly, both in terms of Type I error and power. All three methods failed to correct the population stratification when *F*_*st*_ = 0.001, which is consistent with the failure of the three methods in revealing the admixture structure in the previous experiment. Finally, these results are in line with the results in (14).

**Table 1:** Proportion of associations reported as statistically significant (*P* < 0.0001) by logistic regression using a likelihood ratio test. Random SNPs with no association to the disease were generated by simulating random drift with *F*_*st*_ divergence. Differentiated SNPs with no association to the disease were generated by assuming population allele frequencies of 0.8 of ancestry 1 and 0.2 of ancestry 2. Causal SNPs were generated by combining a multiplicative disease risk model while simulating the random drift with the same *F*_*st*_ as the random SNPs. See methods for more details on the parameters.

|  | Naive | PCA | MDS | SUGIBS |
| --- | --- | --- | --- | --- |
| **$F_{st} = 0.001$** |  |  |  |  |
| **Random** | 0.0002 | 0.0001 | 0.0001 | 0.0001 |
| **Differentiated** | 0.9960 | 0.4483 | 0.6370 | 0.5200 |
| **Causal** | 0.5295 | 0.4779 | 0.4865 | 0.4807 |
| **$F_{st} = 0.005$** |  |  |  |  |
| **Random** | 0.0009 | 0.0001 | 0.0001 | 0.0001 |
| **Differentiated** | 0.9980 | 0.0002 | 0.0003 | 0.0002 |
| **Causal** | 0.5226 | 0.4249 | 0.4255 | 0.4253 |
| **$F_{st} = 0.01$** |  |  |  |  |
| **Random** | 0.0030 | 0.0001 | 0.0001 | 0.0001 |
| **Differentiated** | 0.9971 | 0.0001 | 0.0001 | 0.0001 |
| **Causal** | 0.5166 | 0.4227 | 0.4230 | 0.4229 |

Putting SUGIBS to practice, we projected 2,882 unrelated individuals from a large admixed and heterogeneous dataset containing individuals from varying ancestries (the PSU cohort, see Methods) and eight famous ancient DNA samples onto the first 25 SUGIBS axes established from the 26 populations in the 1KGP. Shown in Figure 5 and S1 (a), the first two ancestry components separate the African (AFR) and East Asian (ESA) populations from the remaining populations, as indicated by the population labels given in the 1KGP. The next two ancestry components in Figure 5 and S1 (b) separate the South Asian (SAS) population and visualizes the admixture in the Admixed American (AMR) population, respectively. In figure 5 and S1 (c), the sixth ancestry component captures different subpopulations in the EAS population. In Figure 5 and S1 (d), the seventh ancestry component is driven by African subpopulations and the separated European subpopulation on the eighth ancestry component is the population from Finland (FIN). The projected PSU cohort is indicated by gray dots in Figure 5 and S1 and overall it is observed that they overlay well with a wide range of ancestry variations in the 1KGP confirming the heterogeneous and admixed nature of the PSU dataset. However, some populations in the 1KGP are less covered by the PSU cohort, such as the population of Finland in Europe and some African subpopulations on ancestry components seven and eight (Figure 5 d).

**Figure 5:**
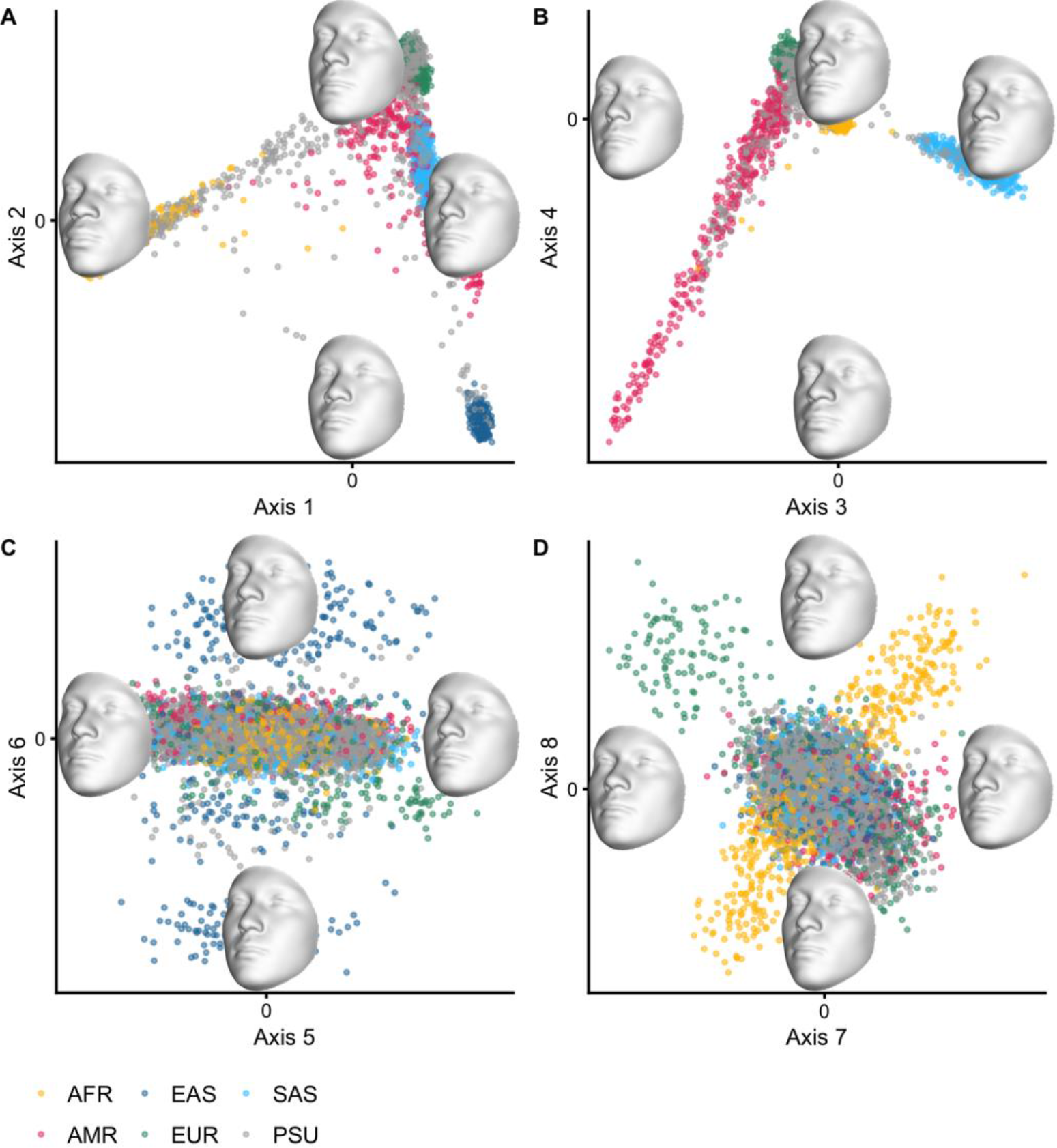
Top eight SUGIBS axes of 1KGP and projections of the PSU cohort. Grouped populations of the 1KGP are coloured dots. The projected PSU cohort are represented by grey dots. The faces illustrate opposing variations along each of the ancestry components and are not associated to any of the 1kG populations in particular (these are shown in Figure 6).

Based on the visually strong and recognizable human facial phenotype, we generated comprehensive illustrations of the population structure embedded in the 1KGP. Using the first 25 SUGIBS scores of the PSU cohort onto the ancestry components of the 1KGP, we fitted a partial least squares regression (PLSR) to model facial variations in function of each of the first eight ancestry components (Figure 5). Strong facial differences are observed for ancestry components 1-4, whilst perceptually smaller differences occur in components 5-8. This is most likely due to a lower overlap of the PSU cohort with these ancestry components. Subsequently, we reconstructed the ancestry population average face from each of the 26 populations in the 1KGP (Figure 6), and ancestry faces specific for eight high-coverage ancient DNA profiles (Figure 7). The facial images in Figures 5, 6 and 7, are perceptually easy to confirm the expected variations in facial shape in function of genetic ancestry including admixtures. For the ancient DNA profiles labeled in Figure 7, it is observed that their projections within the 1kG ancestry is consistent with the geographical locations where these samples were discovered and what is currently known about these samples (Supplementary Table S1).

**Figure 6:**
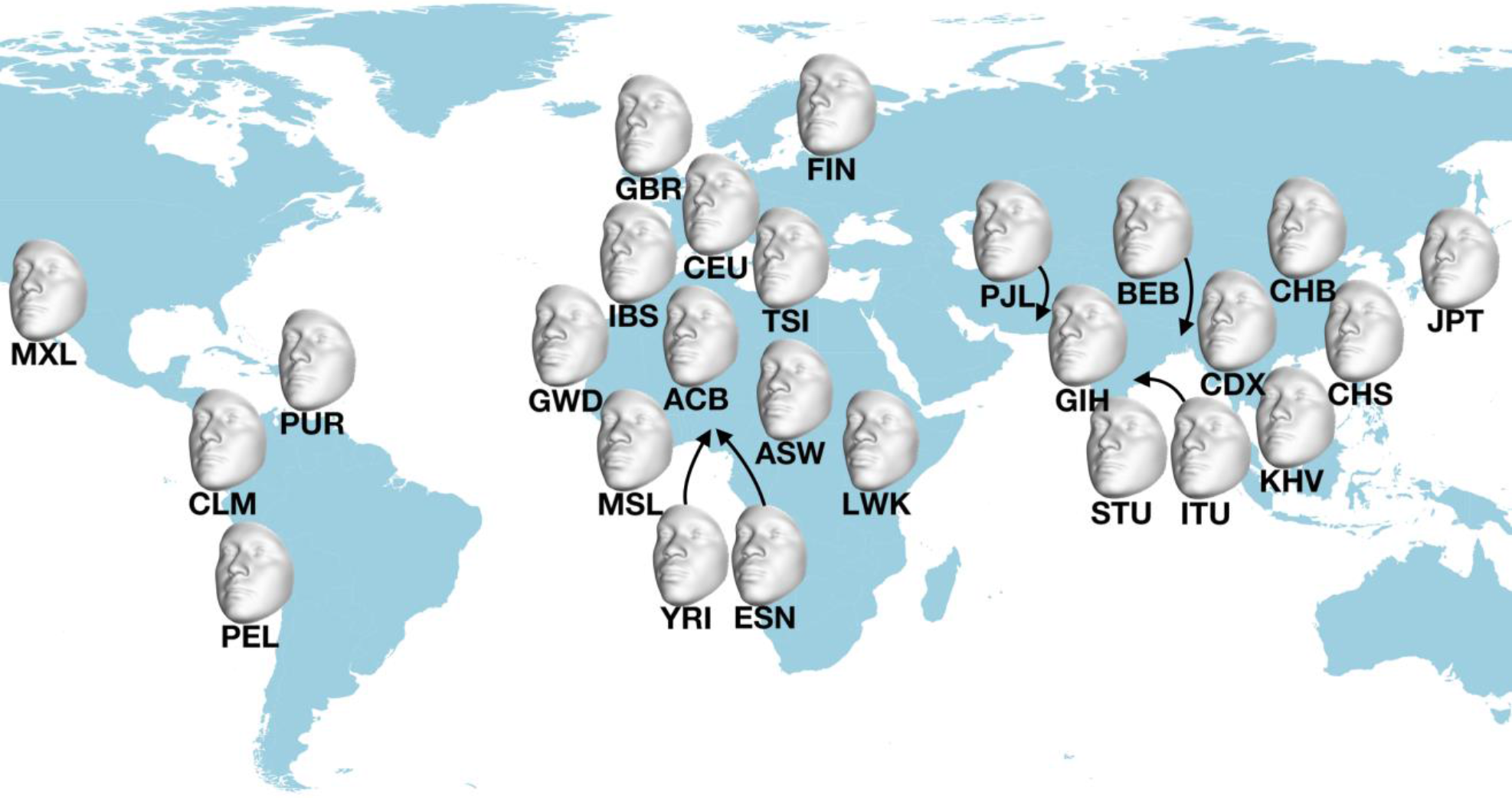
Ancestry population average faces for each of the 26 populations in the 1KGP positioned according to geographical origin. The values for sex, BMI and age in the PLSR model were set to 0 (sexless), 20 and 25, respectively.

**Figure 7:**
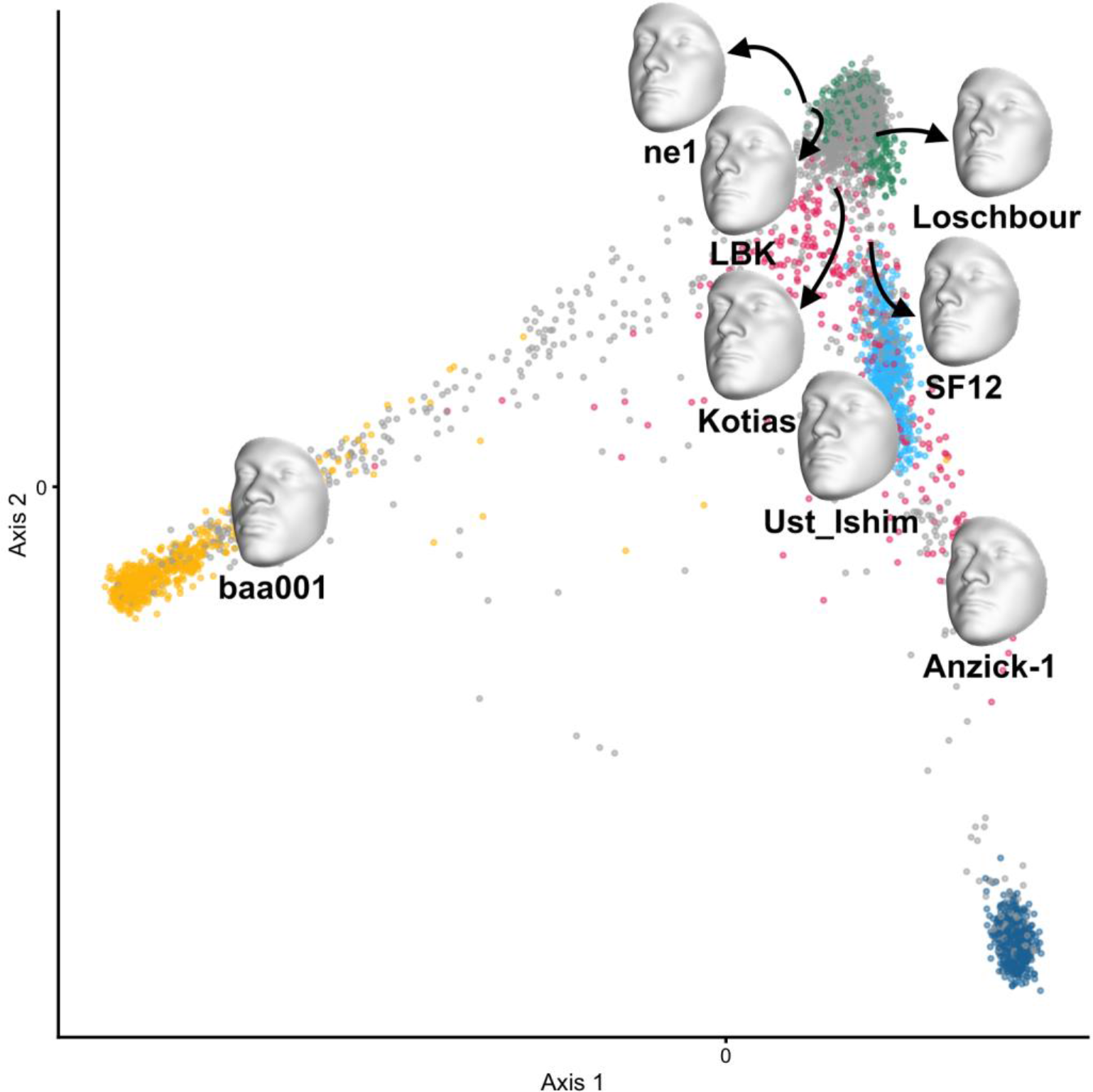
Ancestral facial reconstructions for eight ancient DNA profiles. For these reconstructions, the sex was known from the DNA profile and taken into account in the PLSR model. The values for BMI and age were 20 and 25, respectively.

## Discussion

Accurate inference of population structure and individual global ancestry is of critical importance in human genetics, epidemiology, and related fields (15,16). The analysis of population structure in itself can yield significant insights in terms of population dynamics, both in modern and ancient populations (17–19). Through inspection of ancestry components as well as distances in genetic latent spaces created by, for example, Principal Component Analysis (PCA), it is possible to infer patterns of gene flow and population movements through time. Furthermore, the inclusion of various populations in genome-wide association studies (GWAS) could increase statistical power and make a better contributions to our understanding of the genetics of complex traits for the human population as a whole (20). However, the widely used approach of PCA and analogous techniques are sensitive to outliers, when constructing ancestry spaces, and produce patterns of misalignment due to artifacts of different laboratory protocols when new samples are projected onto a reference ancestry space (1,7,9). We propose a robust alternative for genome-wide ancestry inferencing, referred to as SUGIBS. Our results confirm the erroneous influences in PCA based ancestry estimations that are misleading without careful interpretation. In constructing an ancestry space SUGIBS, shares the same robustness against individual outliers as MDS or related spectral graph approaches (21). Furthermore, and more importantly, during dataset projections SUGIBS is robust against typical artefacts from different laboratory protocols. In addition, SUGIBS achieved the same performance, under error-free conditions, as PCA in revealing the underlying structure of an admixed population and avoiding false positive findings in a simulated case-control GWAS with an admixed population.

Like MDS and SUGIBS, PCA is also a “spectral” method, in which the edge similarity between individuals is simply the covariance of normalized genotypes, commonly referred to as the genomic relationship matrix (22). However, this covariance similarity used in PCA depends on the allele frequencies as a non-robust sample statistic to normalize the genotypes, which causes sensitivity to individual outliers. Note that in our experiments on PCA without using allele frequencies (UPCA) robustness against individual outliers was observed. Among the “spectral” methods, some other robust alternatives were introduced to infer population structure, including a modified genomic relationship (21,23). MDS or related spectral graph approaches (21) using IBS and Allele Sharing Distance (ASD) similarities between individuals (available in PLINK (10)) are also a robust alternative against individual outliers, as illustrated in our results. IBS and ASD are unnormalized distances, and thus less influenced by outliers. However, MDS and the modified genomic relationship used in (21,23), both lack the ability to project new samples on an already established reference ancestry space. Alternatively, it might be possible to use one of the many robust PCA approaches that have been investigated for general data (24–26) as well as genetic data (27). However, in most study data processing protocols, robust approaches are usually used for outlier detection rather than inferring population structure, which is done by classical PCA after excluding outliers (27). This is for example, a standardly used option in the popular EIGENSOFT software (7). Note that, when establishing an ancestry space from a reference dataset, it remains good practice to identify and remove individual outliers, if they are of no further interest.

The main contribution of SUGIBS is robustness against batch artifacts of different laboratory and data processing protocols when projecting new samples onto a reference ancestry space. In the case of missing genotypes, smaller absolute PC scores, and smaller UPC scores are wrongfully generated during the projection of samples. These smaller and decreased scores lead to the “shrinking” and “shifting” patterns as observed in the results. (Note that this is not to be confused with PCA shrinkage due to high dimensional and large-scale data, which is dealt with using shrinkage eigenvalue estimations as recently implemented in EIGENSOFT). However, to correct for this, the projected SUGIBS score matrix is weighted by the reference degree matrix, which captures the similarity between the data to be projected and the reference data (see Methods). This weighting of projected SUGIBS scores equally corrects for the effects of genotyping and imputation errors, as demonstrated in the results. To the best of our knowledge, we are currently not aware of another related approach that offers the same robustness. Based on the results, we argue that SUGIBS is a solid alternative to PCA and MDS and requires less stringent data filters to operate. Our implementation of SUGIBS uses the randomized singular value decomposition algorithm (28), that is also used in FastPCA (12). This makes the algorithm computationally tractable for datasets with tens of thousands of individuals and millions of SNPs. SUGIBS is available as part of an open-source in-house Matlab™library, referred to as <u>SNPLIB</u>, in which we used PLINK binary file formats as input, and provide FastPCA, logistic GWAS and all other methods and simulations mentioned throughout this work. Furthermore, SUGIBS can easily be incorporated into existing and interesting extensions to derive common ancestry estimations in datasets with non-overlapping genetic variants (1), or genotyping-by-sequencing data (29), or population structure inference in presence of relatedness (30), or in iterative schemes to obtain global to fine-scale ancestry estimations (31).

There are a few points of discussion and future investigations. First, a genetic similarity measure between pairs of individuals aims to identify how they are related and different measures exist for ancestry estimations (e.g. IBS, ASD, Identity-by-descent, normalized covariance) (22). Commonly used similarity measures are normalized, just like the traditional approach of PCA on normalized genotype data, to take the genetic composition of individuals along with the rest of the sample into account. A normalization does have the advantage that individuals within the same population are more similar to each other than to individuals in other populations (22). In other words, the distinction between populations increases, which improves population identification by clustering algorithms. However, when the normalization is performed incorrectly clustering efforts might be inaccurate. Furthermore, as seen in our results, such a normalization increases the influence of individual outliers. Finally, in contrast to homogeneous datasets, normalization of genotype data in heterogeneous datasets is challenging depending on whether the dataset is unlabeled or not, imbalanced or not, and with high admixture or not. Starting from unlabeled data, unsupervised clustering approaches such as ADMIXTURE (32) and STRUCTURE (33), iteratively identify the populations individuals belong to and update the normalization accordingly. However, this involves additional parameters to set and tune, the most important one being the amount of clusters expected in the data. Without prior knowledge on how to set these parameters, this can turn into a challenging task. With highly admixture data, any clustering of global ancestry into populations is even questionable. In these situations, only local ancestry estimations, using chromosome painting approaches (34) for example, are meaningful. Alternatively, in the future, we want to investigate the use of a reference ancestry space as constructed in this work, without assigning individuals to specific populations, in estimating normalized genotype data on an individual-by-individual basis. I.e., an ancestry space from unnormalized genotype data is a good first step unbiased by any sample statistics, to further deduct statistics related to individual genotype profiles. For example, (35) propose the Robust Unified Test for Hardy-Weinberg Equilibrium in the context of an admixed population, which also makes use of individual-level adjustments for ancestry. Second, future investigations of the methodology also include the influence of LD pruning and data filtering for SNP selection. Population admixture is one of the main sources for LD between SNPs, therefore we prefer to avoid excessive LD pruning before applying SUGIBS. As stated in (22) any data pruning or filtering is bound to loose information related to population structure. For example, less common variants are typically lost in data filtering, but these might contain valuable information about population structure (22). Since SUGIBS is robust and computationally tractable, any data filtering can be minimized. Third, another future investigation involves the determination of the number of relevant or significant components in SUGIBS, for which we provide a preliminary suggestion that compares the spectrum of the data observed with that of a simulated homogenous dataset assuming linkage equilibrium (LE) and Hardy-Weinberg Equilibrium (Supplementary Text S1).

In application of SUGIBS we used the human face, which is a powerful phenotype to visualize and illustrate underlying genetic ancestry variations. Indeed, faces are easy to recognize, interpret, and validate the outcomes based on everyone’s expert knowledge in facial perception. The faces illustrating the ancestry components of the 1KGP in this work overlay well with the provided population labels. Therefore, they can also provide a means to interpret ancestry variations in a heterogeneous dataset in absence of population labels. It is important to note that an ancestry face, as referred to in this work, for each of the 26 1kG populations and ancient DNA profiles are faces that reflect a population’s or an individual’s genetic background and sex. In other words, ancestry faces are not individually specific faces, but average faces that simply visualize the ancestry background of a DNA profile. Related work on facial prediction from DNA (36,37), also show that sex and ancestry are the primary factors driving the estimation of facial shape from DNA.

Ancestry facial predictions have good value in a range of applications. In archeology, ancestry faces reconstructed from ancient DNA profiles, as done in this work, is of strong interest. Generally, for ancient DNA profiles, missing data is abundantly present, making SUGIBS an interesting technique to be used. Note that, the ancestry faces are limited to modern facial constructs, due to the contemporary facial data used. However, they can help to bring ancient DNA profiles into the context of present-day populations for which facial images (e.g. open-source facial databases, Google images, etc.) are available but DNA is not. Furthermore, there is a good relationship between the face and the skull (38,39), such that ancestry faces can be used to compare against skeletal remains. In the future, it is of interest to deploy our work on datasets of 3D skeletal craniofacial surfaces extracted from Computer Tomography (CT) or Magnetic Resonance Imaging (MRI). In medicine, and more particularly in oral and maxillofacial surgery, the surgical reconstruction of a patient’s face benefits from a proper notion of normal facial shape (40). In the next five to 20 years, whole genome sequencing will become the standard of care in clinics and a patient-specific ancestry face provides a personalized norm of facial shape towards precision medicine in surgical planning. Finally, in forensics, an ancestry facial prediction circumvents the often legally debated reporting of ancestry proportions of a probe DNA profile in a criminal investigation. In France, for example, DNA phenotyping of externally visible traits is legally allowed, since such traits are considered to be public. However, and in contrast, genomic ancestry proportions, as typically reported in forensic DNA testing, is considered to be private information and cannot be used during criminal investigations. We agree that ancestry proportions are not an externally visible characteristic of an individual. The construction of ancestry proportions is also inherently flawed by labelling the individual into so-called parental populations. Furthermore, such numeric information is hard to interpret and use by a forensic investigator. The reconstruction of an ancestry face on the other hand, avoids needing to explicitly label a DNA profile in function of parental populations and provides a visual feedback to an investigator that is perceptually useful, even in admixed cases. A future challenge in forensics does involve the ability to reconstruct ancestry faces using often limited and contaminated DNA material.

In conclusion, SUGIBS is a novel approach to construct an ancestry space from a reference dataset and to project new samples from heterogeneous datasets for a consistent and robust inference of individual ancestry. The main contributions involve robustness against outliers during the construction of an ancestry space, and robustness against batch artefacts during the projection of new samples into an ancestry space. Therefore, SUGIBS is a solid alternative to PCA and MDS and facilitates a robust integrative analysis for population structure and ancestry estimations for heterogeneous datasets. Based on the visually strong and recognizable human facial phenotype, comprehensive illustrations of genomic ancestry variations were provided for different populations in the 1KGP and for eight eminent ancient-DNA profiles. Ancestry facial imaging from genome data has interesting future applications in personalized and precision medicine along with forensic and archeological DNA phenotyping.

## Materials and Methods

### SUGIBS latent-space construction

Given a dataset with *N* individuals and *M* SNPs, we first create an unnormalized genotype (UG) matrix ***X***_***M*×*N***_ with additive genotype coding (*aa* = −1, *Aa* = 0, *AA* = 1 and missing = 0). The UG relationship matrix is then defined as 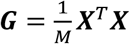. Note that an unnormalized additive genotype coding has only three values (−1, 0, 1) and does not produce extreme values, which occurs with normalized additive genotype encoding schemes (typically used in PCA) due to small minor allele frequencies and in the context of individual outliers.

From ***W***_***N*×*N***_, the IBS similarity matrix of the same dataset used to create ***G***, the similarity degree of an individual can be defined as 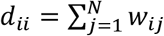. We followed the algorithm implemented in PLINK to calculate the IBS similarity so that:

| IBS | AA | Aa | aa |
| --- | --- | --- | --- |
| AA | 2 | 1 | 0 |
| Aa | 1 | 2 | 1 |
| aa | 0 | 1 | 2 |
| N/A | 0 | 0 | 0 |

However, in contrast to the calculations in PLINK, we do not normalize the IBS similarity matrix with missingness scores. This results in a similarity degree matrix ***D*** defined as the diagonal matrix with *d*_11_, … , *d*_*NN*_ on the diagonal. We use ***D*** to define generalized eigenvectors *ν*_*k*_ = (*ν*_*k*1_, … , *ν*_*kn*_)^𝑇^ of ***G*** with corresponding generalized eigenvalues *λ*_*k*_, and *λ*_1_ ≥ *λ*_2_ ≥ *λ*_3_ ≥ …:

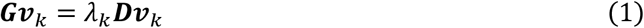

Similar to UPCA, the first generalized eigenvector of ***D*** and ***G*** simply represents the average pattern of all SNPs. Therefore, we start from the second generalized eigenvector and define the *k* th component of SUGIBS to be the *k* + 1th generalized eigenvector of ***G*** and ***D***, *ν*_*k*+1_.

By multiplying 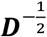 on both sides of equation (1), we obtain:

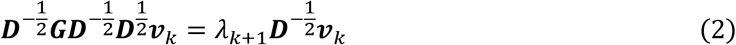

Subsequently, we observe that the eigenvector 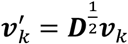 of 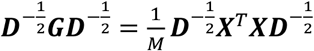 can be obtained from the singular value decomposition (SVD) of the matrix 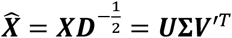, where 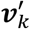 is also the 𝑖th right singular vector with singular value is 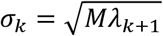, **Σ** is a *N* × *N* diagonal matrix, ***U*** is a *M* × *N* matrix with all the left singular vectors and 𝑽′ is a *N* × *N* matrix with all the right singular vectors.

Denoting ***U***_*k*_ = {***u***_2_, … , ***u***_*k*+1_} and ***Σ***_*k*_ = *diag*{*σ*_2_, … , *σ*_*k*+1_}, the corresponding left singular vectors and the singular values of the first *k* SUGIBS components 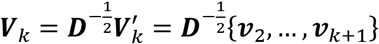, we have the following equation:

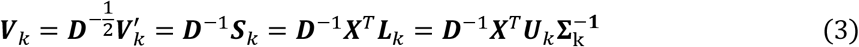

Thus, we denote 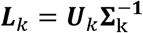 as the SUGIBS loading matrix for the first *k* SUGIBS components and 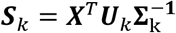 as the unnormalized SUGIBS score matrix.

We proposed a preliminary method to select proper number of components which compared the spectrum of the observed data with that of the simulated data, assuming HWE and Linkage Equilibrium (see Supplement note).

### SUGIBS dataset projection

Given the SUGIBS loadings ***L***_*k*_ from a reference dataset with *N* individuals and *M* SNPs and given a new dataset with 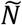 individuals and the same set of SNPs as the reference sample, we denote the unnormalized genotype matrix of the new dataset as 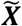. We then define the reference degree 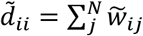, where 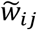 is denoted as the IBS similarity between the *i*th individual in the target dataset and the *j*th individual in the reference dataset. The reference similarity degree matrix 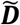 of the new dataset is a diagonal matrix with 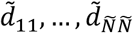on the diagonal. For the first *k* SUGIBS components, the projected score matrix of the target dataset is then obtained as:

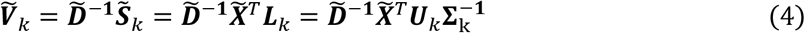

In equation (4), the reference similarity degree matrix 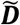 acts as a normalization term correcting the missing genotypes and errors in the samples to be projected. As an example, consider a rare SNP with major allele *A* and minor allele *G*, and an individual with true genotype *AA* that is wrongfully coded as *GG* for that particular SNP. Since the major genotype in the reference data of this SNP is *AA*, the number of shared alleles of this SNP between this individual to the majority of individuals in the reference dataset would reduce from 2 to 0. The unnormalized genotype coding of this person also changes from 1 to −1. Thus, the influence of such a genotyping error on the unnormalized SUGIBS score matrix 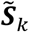 and the reference similarity degree matrix 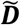 are along the same direction so that the final SUGIBS scores are corrected by 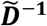. Other typical batch artefact errors and missing genotypes in the new dataset are corrected for in a similar way and, most interestingly, this correction is provided on an individual by individual basis.

### Genome-wide common SNP selection across datasets

We recommend the following procedure to extract a common set of SNPs between a reference dataset and another dataset being projected, for constructing SUGIBS ancestry spaces. First, we exclude all the indel, monomorphic, and multi-allelic SNPs in both the reference dataset and the dataset to project. Subsequently, we extract the list of SNPs common in both datasets. Based on this list, we further recommend a minor allele frequency (MAF) filtering with a MAF threshold of 0.01 on the reference dataset using PLINK (10) as a quality control step. We do not recommend Hardy-Weinberg disequilibrium (HWD) filtering since it is probably the result of population admixture and thus useful for our purposes (41). Although population admixture is one of the main sources for LD between SNPs, we still recommend LD pruning since it is not unusual to have non-uniformly genotyped genomes. Similar to PCA, SUGIBS do not explicitly model LD between SNPs so that misleading results might be generated without LD pruning.

### Individual outlier robustness

The basic dataset that was used to investigate robustness against individual outliers in a reference dataset, consists of the individuals from the CEU population (111 individuals) and the TSI population (102 individuals) from the HapMap 3 dataset (3), after excluding non-founders. We randomly selected one individual as outlier from four other populations (CHB, MEX, GIH, and YRI). These individuals specifically are NA18798 (CHB), NA19740 (MEX), NA21124 (GIH), and NA19262 (YRI). After removing the monomorphic SNPs in each of these three datasets, we built SUGIBS, MDS, UPCA and PCA spaces using 892,338 autosomal SNPs remaining in all three datasets. We intentionally did not perform either minor allele frequency (MAF) filtering or HWE filtering on the SNPs since many rare SNPs and SNPs violating HWE are due to the outliers and were therefore not checked for during the testing for robustness.

### Simulated laboratory artefacts

We used the 1000 Genomes Project dataset (2,504 unrelated individuals from 26 populations) as the reference dataset to infer a PCA, UPCA and SUGIBS based ancestry space. We used the HGDP dataset that analyzed genomic data from 1,043 individuals from around the world as the dataset to project. First, we remapped the HGDP dataset from the NCBI36 (hg18) assembly to the GRCh37 (hg19) assembly using the NCBI Genome Remapping Service. Based on the SNP selection procedure for SUGIBS as explained previously, we further performed a LD pruning with a window size of 50, a moving step of 5 and a threshold *r*^2^ > 0.2 for several times until no more SNPS were excluded, following (12). LD pruning is a common practice when using PCA. Therefore, we followed this additional step to make the results based on PCA, UPCA and SUGIBS comparable. We finally selected 154,199 autosomal SNPs to construct the PCA, UPCA and SUGIBS ancestry spaces. We then extracted the first eight PCA, UPCA and SUGIBS ancestry components from the reference dataset. After extracting the same set of SNPs in the HGDP dataset, we took care to ensure that the alternate alleles were the same as in the reference dataset.

Since PLINK binary file format stores the genotypes of four consecutive individuals in a single byte, we assigned one of every two “bytes” (four individuals) into Population A and the other individuals into Population B of the HGDP dataset. This resulted in 523 individuals for Population A and 520 individuals for Population B. In order to simulate laboratory artefacts, we randomly masked 5% genotype calls as missing and changed 5% genotype calls (e.g., from AA to Aa or aa) of the rare SNPs (MAF < 0.05) in Population A. Random genotype masking and changing were also performed on the “byte” level, i.e. four individuals at a time. For both genotyping masking and changing, we generated 100 datasets to project on the 1kG reference ancestry space. Subsequently, we calculated the root-mean-square deviations (RMSD) between the scores of the top eight PCA, UPCA and SUGIBS axes generated using the original genotypes and the modified genotypes in Population A and further normalized them by the range of the axes generated using the original genotypes so that normalized root-mean-square deviations (NRMSD) across methods are comparable.

### Simulated admixed population

Our admixture simulations were adapted from the section “Simulation Framework” in (12). For a given SNP *i*, the ancestral allele frequency *p*_*i*_ was sampled from a *uniform* (0.1,0.9) distribution. Population allele frequencies were generated by simulating random drift in two populations of fixed effective size *N*_*e*_ for *τ* generations as *p*_*i*1_ and *p*_*i*2_, whose initial values were set to *p*_*i*_. In each generation, the number of alternate alleles *Z*_*i*1_ and *Z*_*i*2_ were sampled from two binomial distributions with 2*N*_*e*_ number of trials and *p*_*i*1_ and *p*_*i*2_ success probabilities. The population allele frequencies were then updated by 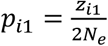 and 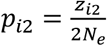 For all simulations, population allele frequency simulations were run for 20 generations and the effective population size *N*_*e*_ was calculated for a target 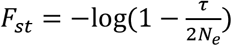(42). This was done for *F*_*st*_ = {0.001,0.005,0.01,0.05,0.1}, *N*_*e*_ ≈ {10*k*, 2*k*, 1*k*, 200,100} with *τ* = 20.

The ancestry proportions *α*_*j*_ were sampled from a *beta* (0.5, 0.5) distribution so that the proportion from each ancestry is 50% on average. For a given individual *j* with ancestry proportion of *α*_*j*_ from Population one and (1 − *α*_*j*_) from Population two, the individual allele frequency for SNP *i* was 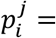 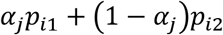 and the genotype was sampled from a binomial distribution with 2 trials and 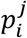success probability. The Matlab™ implementations for these simulations are also provided in our SNPLIB library.

### Simulated GWAS

Our GWAS simulation is similar to the one carried out in (14). To simulate a case-control GWAS, we generated 1,000 individuals from a population admixed from two ancestries. The case-control status was simulated using a disease risk proportional to *r*^*α*^, based on an ancestral risk of *r* = 3. We generated three categories of SNPs (random, differentiating and causal) to compare the performance of PCA, MDS, and SUGIBS in correcting for population stratification. For the first category (random SNPs with no association to the disease), we generated the SNPs by simulating random drift with a certain *F*_*st*_ divergence. For the second category (differentiated SNPs with no association), we assumed population allele frequencies of 0.8 for ancestry one and 0.2 for ancestry two. For the third category (causal SNPs), we generated SNPs by combining a multiplicative disease risk model while simulating the random drift with the same *F*_*st*_ as the random SNPs.

We simulated the case-control status according to (7). For individuals with an ancestry proportion of *α* from population one and (1 − *α*) from population two, the case-control status was simulated with the probability of disease equal to 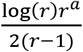, which ensures an average value of 0.5 across all the values of *α* (7).

For the case individuals, the population allele frequencies *p*_*i*1_ and *p*_*i*2_ of the causal SNP *i* were further 529 updated to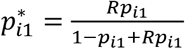and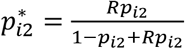 with a relative risk of *R* = 3, respectively. The Matlab™ implementations for these simulations are also provided in our SNPLIB library.

### PSU cohort and 3D facial images

Study participants in the PSU cohort were recruited in the United States through several studies based at The Pennsylvania State University under Institutional Review Board (IRB) approved protocols (IRB #44929, #45727, #2503, #4320, #32341). 3D facial images were taken using the 3dMD Face (3dMD, Atlanta, GA) and the Vectra H1 (Canfield, Parsippany, NJ) imaging systems. Height and weight were measured using an Accustat stadiometer (Genentech, San Francisco, CA), a clinical scale (Tanita, Arlington Heights, IL), or by self-report. Genotyping was conducted by 23andMe (23andMe, Mountain View, CA) on the v4 genome-wide SNP array and on the Illumina Multi-Ethnic Global Array (MEGA). After filtering out SNPs with more than 10% missing genotypes, the intersection of these two arrays compromised of approximately 600K SNPs. We removed individuals with misclassified sex information, missing covariate data, and those with more than 10% missing genotypes. Relatives were identified as pairs of individuals with an identity-by-state (IBS) value of at least 0.8, after which one of each pair was randomly removed, resulting in a set of 2,882 individuals. Genotypes were imputed to the 1000 Genomes Project Phase 3 reference panel, using SHAPEIT2 (Delaneau, Marchini, & Zagury, 2012) for prephasing of haplotypes and imputed using the Sanger Imputation Server PBWT pipeline (Durbin, 2014; McCarthy et al., 2016).

3D facial images were imported into Matlab™ 2016b in .obj wavefront format to perform spatially dense registration (MeshMonk). After importing the images, five positioning landmarks were indicated in the corners of the eye, the tip of the nose and the corners of the mouth to roughly align the images into the same position. Subsequently, the images were cleaned by removing hair, ears, and any dissociated polygons. A symmetrical anthropometric mask (43) of 7,160 landmarks was then mapped onto the pre-processed images (44). This resulted in homologous spatially dense configurations of quasi-landmarks per facial image. Reflected images were created by changing the sign of the x-coordinate of the original mapped images. Both the original and the reflected remapped faces were then superimposed following a generalized Procrustes superimposition to eliminate differences in orientation, position and scale (45). Symmetrized images were created by averaging the original and the reflected images.

Image quality control was performed to identify poorly remapped faces using two approaches. First, as described in (46), outlier faces were identified by calculating Z-scores from the Mahalanobis distance between the mean face and each individual face. Faces with Z-scores higher than 2 were manually checked. Second, a score was calculated that reflects the missing data present in the image due to holes, spikes, and other mesh artefacts that can be caused by facial hair or errors during the pre-processing steps, for example. Images with scores indicating a high amount of missing data, indicating large gaps in the mesh, were also manually checked. During the manual check, the images were either classified as images of poor quality or were pre-processed again if possible and mapped again.

### Prediction of ancestry faces

Using 69,194 autosomal SNPs overlapping with the PSU cohort and the ancient-DNA profiles, we constructed 25 SUGIBS ancestry components, which is theoretically sufficient to separate 26 populations, from the 1000 Genomes project. Subsequently, we projected the individuals from the PSU cohort and the ancient-DNA profiles onto the 1kG ancestry components. Then, we fitted a partial least-squares regression (PLSR) model using the superimposed 3D facial images with 7,160 quasi-landmarks collected in the PSU cohort as the response variables and the 25 projected SUGIBS scores of the PSU cohort together with three covariates (age, sex, and BMI) as the explanatory variables.

Given specific ancestry scores on the ancestry components of the 1kG ancestry space, together with age, BMI and sex (−1 (male), 0 (neutral sex) or 1 (female)), the PLSR model was used to predict ancestry faces. To illustrate the ancestry components in Figure 7, we simply varied a single score along each ancestry component separately, while keeping the scores on the other ancestry components fixed and equal to the overall average scores in the PSU cohort together with values for age = 25, BMI = 20, and sex = 0. For each of the 26 populations in the 1KGP, we calculated the average scores on each SUGIBS ancestry component per population. These average scores together with values for age = 25, BMI = 20, and sex = 0, were used in the PLSR model to reconstruct the average ancestry faces for each of the 26 populations in the 1KGP. Diploid genotypes for the ancient genomes were called using GATK as described in (47). The projected scores of the ancient-DNA profiles were used together with the genome-derived sex values of each of the ancient individuals to reconstruct their ancestry faces in Figure 7.

## Acknowledgments

### Funding

Jiarui Li is in part supported by a Chinese Research PhD scholarship. This investigation was also supported by the KU Leuven, BOF, FWO Flanders (G078518N) and the NIH (1-RO1-DE027023). Kristel Van Steen acknowledges research opportunities offered by F.N.R.S (Convention n° T.0180.13) and by the interuniversity research institute Walloon Excellence in Lifesciences and BIOtechnology (WELBIO-Convention de Recherche n° WELBIO-CR-2015S-03R). The collaborators at the Penn State University were supported in part by grants from the Center for Human Evolution and Development at Penn State, the Science Foundation of Ireland Walton Fellowship (04.W4/B643), the United States National Institute Justice (www.nij.gov; 2008-DN-BX-K125), and by the United States Department of Defense (www.defense.gov). Torsten Günther is supported by the Swedish Research Council (2017-05267).

### Ethics Statement

Institutional review board (IRB) approval was obtained at each recruitment site and all participants gave their written informed consent prior to participation; for children, written consent was obtained from a parent or legal guardian. For the PSU cohort, the following local ethics approvals were obtained: State College, PA (IRB #44929 and #4320 New York, NY (IRB #45727); Urbana-Champaign, IL (IRB #13103); Dublin, Ireland; Rome, Italy; Warsaw, Poland; and Porto, Portugal (IRB #32341); and Twinsburg, OH (IRB #2503)

### Author contributions

J.L under supervision of P.C and K.V.S developed the SUGIBS methodology. J.L. together with P.C. designed the experiments with input from K.V.S and M.D.S. J.L. under supervision of P.C. and M.D.S conceptualized and implemented the ancestral facial imaging based on the 1000 Genome Project. J.W., T.G., A.Z., R.J.E. and S.W. curated the genomic data, including that of the PSU cohort. M.D.S, J.W., K.I., H.H., N.N., and A.O.C collected and processed the 3D facial image data of the PSU cohort. T.G., E.M.S., and M.J., provided and curated the eight ancient DNA profiles and were involved in the ancestry facial imaging thereof. J.L and P.C wrote the manuscript with extensive input from all co-authors.

### Data and materials availability

Most data used in this work originates from public open-source projects, including the HapMap 3 project, 1000 Genome project and the HGDP dataset. For access to this data, we refer to their respective webpages as indicated under the URL section.

The participants comprising the Penn State University dataset (PSU cohort) were not collected with broad data sharing consent. Given the highly identifiable nature of both facial and genomic information and unresolved issues regarding risk to participants, we opted for a more conservative approach to participant recruitment. Broad data sharing of these collections would thus be in legal and ethical violation of the informed consent obtained from the participants. This restriction is not because of any personal or commercial interests. Additional details and a more confined sharing can be requested from M.D.S.

An implementation of SUGIBS is freely available (see URL Section). This comprises a Matlab™ toolbox, referred to as SNPLIB, and contains implementations of all the methods and simulations used in this work. We also provide the resulting PLSR model, with demo script, to create Ancestry Facial images for other open or in-house data collections (currently under construction). The spatially-dense facial mapping software, referred to as MeshMonk, is available free of use for academic purposes (see URL Section).

### URL’s

HapMap 3 Data: https://www.genome.gov/10001688/international-hapmap-project/

1000 Genome Project: http://www.internationalgenome.org/

HGDP dataset: http://www.cephb.fr/hgdp/

SNPLIB: https://github.com/jiarui-li/SNPLIB

MeshMonk: https://github.com/TheWebMonks/meshmonk

NCBI Genome Remapping Service: https://www.ncbi.nlm.nih.gov/genome/tools/remap

## Supplementary Materials

**Supplementary Table S1:** Information and references for each of the 8 ancient DNA profiles.

**Supplementary Figure S1:** Top eight SUGIBS axes of 1KGP and projections of the PSU cohort

**Supplementary Text S1:** Determination of the number of relevant or significant components

