## Supplementary Materials for "Robust Genome-Wide Ancestry Inference for Heterogeneous Datasets and Ancestry Facial Imaging based on the 1000 Genomes Project"

### Supplementary Table S1:

Table S1: Information and reference on eight ancient DNA profiles.

| Individual ID | Geography | Time (years cal BP) | Reference |
|---|---|---|---|
| Anzick-1 | North America | 12,707–12,556 | Rasmussen et al. (2014)(1) |
| baa001 | Southern Africa | 1,986–1,831 | Schlebush et al. (2017)(2) |
| Kotias | Caucasus | 9,529–9,895 | Jones et al.(2015)(3) |
| LBK | Europe | 7,450-6,750 | Lazaridis et al. (2014)(4) |
| Loschbour | Europe | 8,170-7,940 | Lazaridis et al. (2014)(4) |
| ne1 | Europe | 7,020–7,290 | Gamba et al. (2014)(5) |
| SF12 | Northern Europe | 9,033–8,757 | Günther et al. (2018)(6) |
| Ust_Ishim | Western Siberia | 46,880–43,210 | Fu et al. (2014)(7) |

**Supplementary Figure S1:**

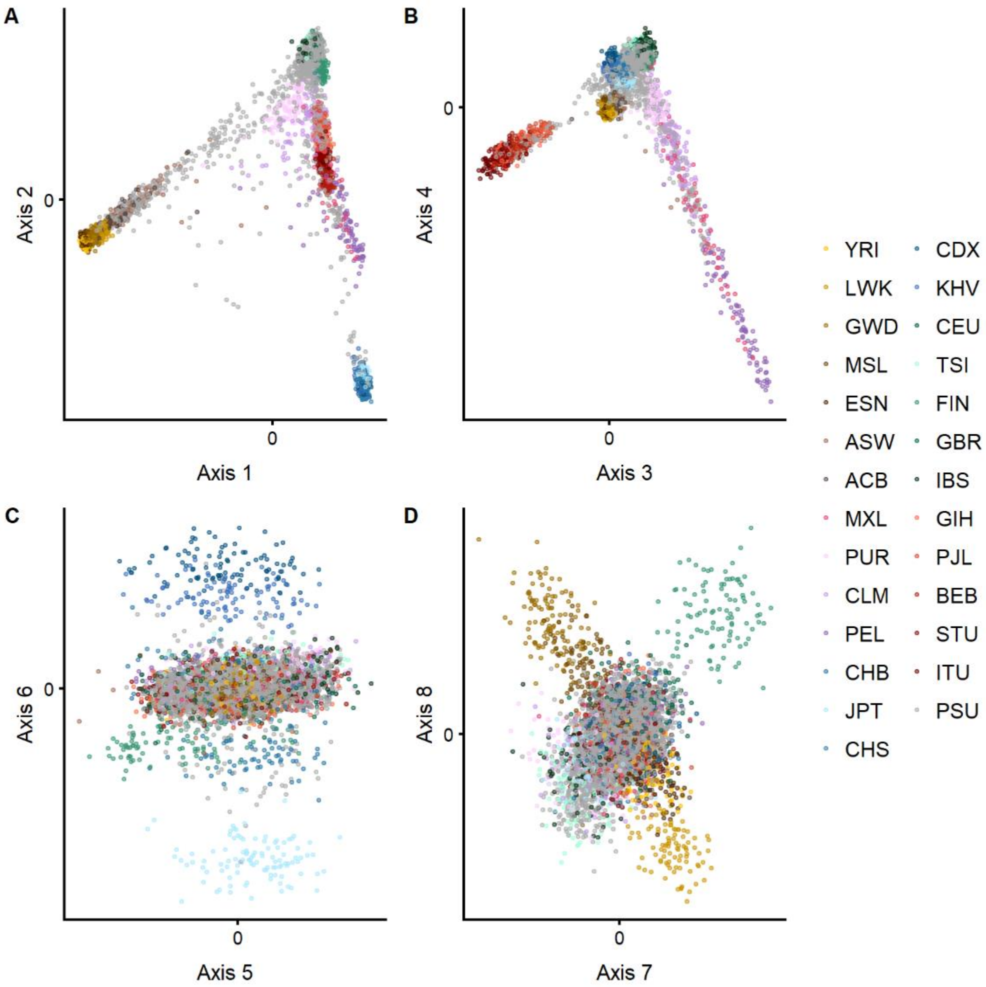


*Figure S1: Top eight SUGIBS axes of 1KGP and projections of the PSU cohort. Grouped populations of the 1KGP are coloured dots. The projected PSU cohort are represented by grey dots. In contrast to Figure 5, the populations are not grouped into the larger continental population groups.*

**Supplementary Text S1:** Determination of the number of relevant SUGIBS components

A key question for any lower dimensional embedding of data into a latent-space is the determination of the number of relevant latent components. In previous work (8), we used PCA to obtain lower dimensional facial shape presentations in combination with a technique referred to as Parallel Analysis (9,10). A Parallel Analysis determines the amount of eigenvalues (and thus the number of principal components (PCs)) from the observed data that are significantly different from eigenvalues computed from permuted versions of the original data. By running multiple permutations, a null distribution of noisy eigenvalues is obtained, against which significance of the original eigenvalues can be tested (whilst taking the properties of the data itself into account). Similar to a Parallel Analysis in PCA(9), our preliminary method or suggestion to select the number of components for SUGIBS is by comparing the spectrum of eigenvalues from an observed potentially heterogeneous dataset (HED) with that of simulated homogeneous datasets (HOD). This is done using the same number of SNPs and samples as in the observed dataset.

For the HODs, we generate the genotypes of each SNP independently according to the allele frequency calculated from the observed data. This implies that each SNP is in HWE but is not in LD with any other SNP. For each simulated HOD and the HED, we calculate the eigenvalues of $\boldsymbol{D}^{-\frac{1}{2}}\boldsymbol{G}\boldsymbol{D}^{-\frac{1}{2}}$, where the unnormalized genomic relationship matrix is defined as $\boldsymbol{G}$ and $\boldsymbol{D}$ is a similarity degree matrix defined by the IBS similarity. By comparing the eigenvalues of the HEDs with the eigenvalues from the simulated HODs, an indication whether the observed dataset deviates from a single homogeneous population is provided. However, we observed that the LD between the SNPs in a sample does affect the sloop of the eigenvalue spectrum. To illustrate this, we simulated three datasets each with 10,000 SNPs and 1,000 samples assuming homogeneity, but with different levels of LD between SNPs (no LD, $r^2 \leq 0.2$ and $r^2 \leq 0.8$). The results in Figure S2 show that the higher the LD level, the steeper the eigenvalue spectrum becomes. In other words, the first eigenvalues explain more of the total variance due to correlation in the data, which is expected given the increased levels of LD.

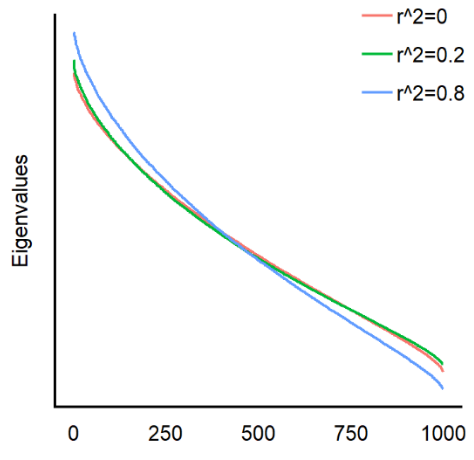


*Figure S2: Spectrum in descending order in function of LD level. Y-axis represents the values of eigenvalues.*

In order to adjust for the different slopes of the eigenvalue spectrum caused by different levels of LD, we fit a robust regression (robustfit in MATLAB) between the observed eigenvalue spectrum and the simulated eigenvalue spectrum. A robust regression, was chosen since it is not influenced by the first few large eigenvalues, which are expected for highly heterogeneous population samples. In practice, we run the simulation 100 times and robustly fit the observed eigenvalue spectrum with the median of the 100 simulated eigenvalue spectrums. Subsequently, we plot the observed eigenvalue spectrum against the adjusted simulated eigenvalue spectrums.

Results for simulated heterogeneous population samples with an admixture from three, six and nine different ancestries with different levels of $F_{st}$ (0.1, 0.01 and 0.001) are shown in Figure S3. It is observed that the simulated HOD eigenvalue spectrum is consistently lower than the observed HED eigenvalue spectrum, and this for all 30 eigenvalues plotted. Therefore, in contrast to Parallel Analysis, the simulated HOD eigenvalue spectrum could not be used as a direct indicator for the number of significant components, since all the observed eigenvalues remain larger (and thus significant) compared to the simulated ones. However, an indication of the amount of relevant (instead of significant) components that represent admixture is still observed. For larger values of $F_{st}$ (0.1, 0.01), the correct number of relevant components (2 for three ancestries, 5 for six ancestries and 8 for nine ancestries), are visually distinct in magnitude in comparison to the simulated HOD eigenvalue spectrum, and this distinction is larger than the subsequent (non-relevant) components. For lower values of $F_{st}$ (0.001), and an admixture from more than 3 ancestries, this visual distinction is lost.

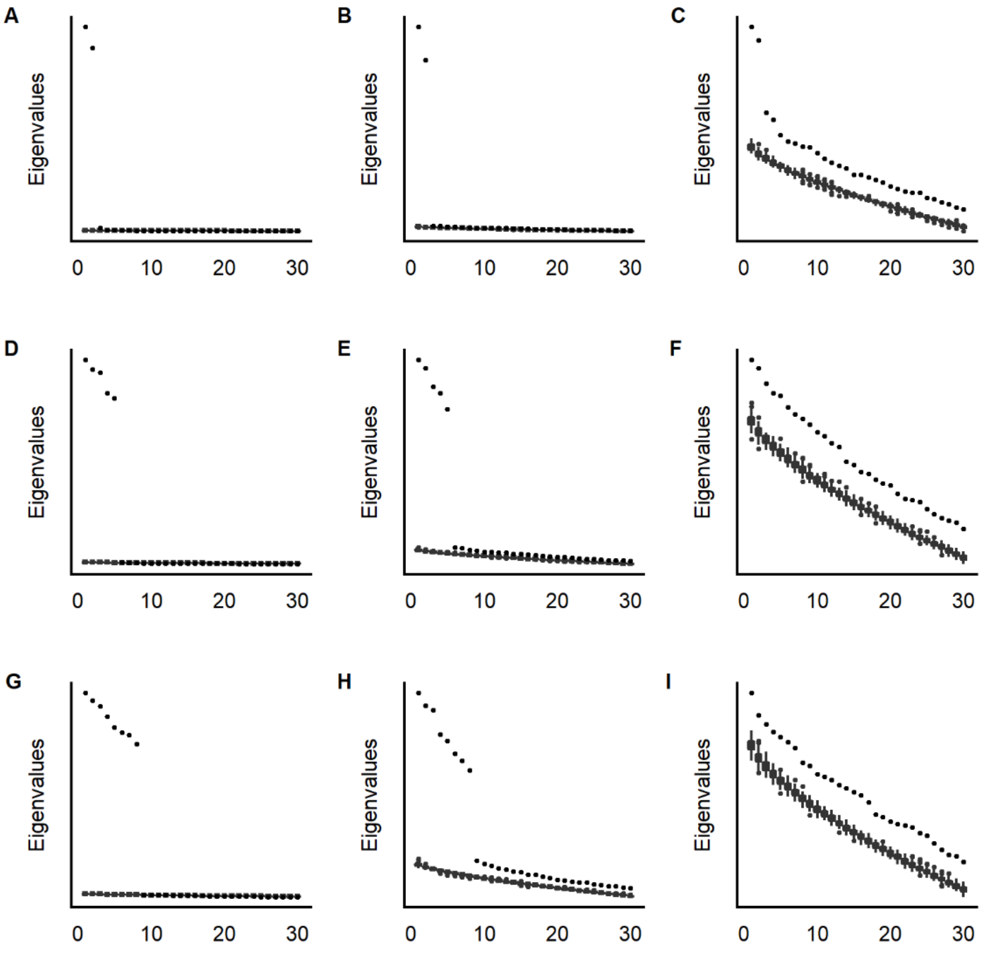


*Figure S3: Indicative evaluation of the number of relevant components for simulated heterogeneous population samples. Each simulated heterogeneous population sample is an admixture from A) three ancestries with* $F_{st} = 0.1$*, B) three ancestries with* $F_{st} = 0.01$*, C) three ancestries with* $F_{st} = 0.001$*, D) six ancestries with* $F_{st} = 0.1$*, E) six ancestries with* $F_{st} = 0.01$*, F) six ancestries with* $F_{st} = 0.001$*, G) nine ancestries with* $F_{st} = 0.1$*, H) nine ancestries with* $F_{st} = 0.01$*, and I) nine ancestries with* $F_{st} = 0.001$*. For each simulated admixed as well as homogenous population sample was generated using 1,000 samples with 10,000 SNPs.*
